# Tau seeding activity anticipates phospho-tau pathology in Alzheimer’s disease

**DOI:** 10.1101/267724

**Authors:** Sarah K. Kaufman, Kelly Del Tredici, Talitha L. Thomas, Heiko Braak, Marc I. Diamond

**Affiliations:** Center for Alzheimer’s and Neurodegenerative Diseases, Peter O’Donnell Jr. Brain Institute, University of Texas Southwestern Medical Center, Dallas, TX, USA; Graduate Program in Neuroscience, Washington University in St. Louis, St. Louis, MO; Clinical Neuroanatomy Section/Department of Neurology, Center for Biomedical Research, University of Ulm, Ulm, Germany

## Abstract

Alzheimer’s disease (AD) is characterized by accumulation of tau neurofibrillary tangles (NFTs) and, according to the prion model, transcellular propagation of pathological “seeds” may underlie its progression. Staging of NFT pathology with phospho-tau antibody is useful to classify AD and primary age-related tauopathy (PART) cases. The locus coeruleus (LC) shows the earliest phospho-tau signal, whereas other studies suggest that pathology begins in the transentorhinal/entorhinal cortices (TRE/EC). The relationship of tau seeding activity, phospho-tau pathology, and progression of neurodegeneration remains obscure. Consequently, we employed an established cellular biosensor assay to quantify tau seeding activity in fixed human tissue, in parallel with AT8 phospho-tau staining of immediately adjacent sections. We studied four brain regions from each of n=247 individuals across a range of disease stages. We detected the earliest and most robust seeding activity in the TRE/EC. The LC did not uniformly exhibit seeding activity until later NFT stages. We also detected seeding activity in the first temporal gyrus and visual cortex at stages before NFTs and/or AT8-immunopositivity were detectable. AD and putative PART cases exhibited similar patterns of seeding activity that anticipated histopathology across all NFT stages. Our findings are consistent with the prion model and suggest that pathological seeding activity begins in the TRE/EC rather than in the LC, and may offer an important addition to classical histopathology.

## Introduction

Tauopathies constitute a diverse group of neurodegenerative diseases that include Alzheimer’s disease (AD). They are defined by the deposition of aggregated phospho-tau protein in the central nervous system^1,2^. Tau aggregation is directly linked to the pathogenesis of tauopathies, as tau mutations that increase the propensity of tau to aggregate cause dominantly inherited dementia^3^. The neuropathology of AD, the most common form of dementia, features intraneuronal pretangle and neurofibrillary tangle (NFT) tau pathology as well as extraneuronal ghost tangles and, in advanced NFT stages, various forms of extracellular amyloid beta (Aβ) plaques. This pathology has a characteristic regional pattern of progression, thereby permitting the distinction of different stages in asymptomatic and symptomatic individuals^4,5^. Recently, it was proposed that early NFT stages with pathological changes confined to the anteromedial temporal cortex, and minimal or no Aβ deposits, may constitute a primary age-related tauopathy (PART)^6^, a hypothesis that remains a source of debate (Duyckaerts et al. 2015). The spatiotemporal pattern of tau pathology in AD correlates well with brain atrophy and cognitive decline observed in patients^4,7^. Based on extensive experimental data, we and others have proposed that transcellular propagation of tau protein “seeds,” in the manner of prions, could underlie the inexorable spread of pathology in tauopathies^8^.

Tau aggregates that accumulate in tauopathies exhibit a high degree of phosphorylation^1^. Traditional immunohistochemistry (IHC) has been the gold standard for disease staging and discrimination among tauopathy syndromes^9–11^. The monoclonal antibody AT8, which recognizes phospho-serine 202 and phospho-threonine 205 on aggregated tau protein, is a principal tool to define AD intraneuronal pathology^12^. The AT8 signal increases with disease progression (Suppl. Fig. 1A-C)^4^. It first appears in the locus coeruleus (LC), and thereafter in a few additional brainstem nuclei with diffuse cortical projections (subcortical stages 1a-c). The first cortical lesions have been observed in neuronal processes (cortical pretangle stage 1a) and in projection neurons (cortical pretangle stage 1b) of the transentorhinal region (TRE) in the absence of Aβ deposits^13^. This led to the idea that tau aggregation in the LC may represent the earliest phase of AD pathogenesis^13,14^. At NFT stage I, Gallyas silver staining reveals neurofibrillary lesions restricted to selected brainstem nuclei and the TRE. Pathology then develops in the entorhinal cortex (EC) of the parahippocampal gyrus at NFT stage II. At NFT stage III, it begins to involve the CA1 sector of the hippocampal formation and enters the neocortical regions of the temporal neocortex adjoining the TRE. NFT stages IV and V are characterized by increasingly abundant tau pathology in neocortical regions. The first temporal gyrus becomes involved at NFT stage V, and during NFT stage VI the primary neocortical areas, such as the primary visual field, exhibit tau lesions^5,7^ (Suppl. Table 1). In comparisons of pathology and clinical presentation, over half of the patients at NFT stages III-IV exhibited signs of mild cognitive impairment, and over 90% of patients at NFT stages V-VI exhibited moderate to severe dementia^9^.

The progressive accumulation of neurofibrillary tau pathology has long been recognized to involve neural networks^4,15^. Recent work *in vitro* and *in vivo*^17–20^ indicates that in experimental systems tau assemblies (seeds) spread pathology between interconnected neurons and progressively trigger further aggregation of native tau. This is similar to the pathophysiology of prion diseases, where prion protein (PrP) adopts a beta sheet-rich conformation that self-assembles and acts as a template to convert native PrP to a pathogenic form^21,22^. In general, transcellular propagation of aggregation appears to be a common feature of various proteins implicated in neurodegenerative diseases^16^,^23–26^.

The term “prion” is controversial as applied to noninfectious neurodegenerative diseases^27–31^. We favor a definition that encompasses the myriad of proteins that exist as monomers or as self-replicating assemblies, and which specify biological activity based on their conformation^8^. Based on the prion hypothesis, we have hypothesized that tau seeding activity will mark incipient, submicroscopic protein aggregation before the occurrence of tau pathology that is visible by light microscopy.

We have previously developed a sensitive and specific cell-based “biosensor” assay to detect tau seeding activity in biological samples^32,33^. When we used this assay in a transgenic mouse model of tauopathy, we observed seeding activity far in advance of detectable histopathology or accumulation of insoluble tau protein^32^. In fresh frozen tissue from AD patients, we have also observed seeding activity in advance of predicted neuropathological changes^34^. However, in such studies fresh frozen samples are more difficult to obtain than fixed brain tissue, and do not allow direct anatomical comparison of seeding activity with high quality histopathology. To resolve this problem, we recently developed a method to quantify tau seeding activity in fixed, archived human brain sections^35^. This has allowed simultaneous IHC and measurement of seeding activity in fixed tissues classified as AD and PART, and in asymptomatic individuals. We have now assessed the relationship of seeding to phospho-tau pathology in the LC and in more distant cortical regions, thereby addressing fundamental questions about AD pathogenesis.

## Methods

### Culture of biosensor cells

Seeding assays were performed with a previously published biosensor cell line that stably express tau-RD(P301S)-CFP and tau-RD(P301S)-YFP (ATCC CRL-3275)^32^. All HEK293 cells were grown in complete media: Dulbecco’s Modified Eagle’s Medium (DMEM) (Gibco) with 10% fetal bovine serum (Sigma) and 1% penicillin/streptomycin (Gibco). Cells were cultured and passaged at 37°C, 5% CO_2_, in a humidified incubator. Dulbecco’s phosphate buffered saline (Life Technologies) was used for washing the cells prior to harvesting with 0.05% Trypsin-EDTA (Life Technologies).

### Tau KO mouse breeding

To determine a true negative tissue for assays, we used tau knockout mice containing a GFP-encoding cDNA integrated into exon 1 of the MAPT gene. These were obtained from the Jackson Laboratory and maintained on a C57BL/6J background^36^. Animals were housed on a 12h light/dark cycle and provided with food and water ad libitum. All animal maintenance and experiments adhered to the University of Texas Southwestern animal care and use protocol.

### Mouse sample collection and preparation

Animals were anesthetized with isoflurane and perfused with chilled PBS with 0.03% heparin. Whole-brains were drop-fixed in 4% paraformaldehyde in PBS overnight at 4°C. Brains were incubated in 30% sucrose before sectioning. Sections were collected to equivalent volume of human samples (100 μm thickness × 4 mm circular punch biopsy) and placed in TBS with protease inhibitors (Sigma Aldrich complete protease inhibitor, EDTA free) as described below. Mouse and human samples were subsequently prepared in an identical fashion.

### Human sample staging and preparation

Human autopsy tissue used for this study was obtained from n=247 individuals with AT8-positive tau pathology (116 females, 131 males, age range 14-97 years, Table 1) and 6 controls (4 females, 2 males, age range 45-72 years) in compliance with ethics committee guidelines at the University of Ulm and the University of Frankfurt as well as German federal and state law governing human tissue usage. The brain specimens included cases from affiliated university hospitals in Germany. The brains were fixed in a 4% buffered aqueous solution of formaldehyde and subsequently archived for up to 25 years. Tissue blocks were excised and embedded in polyethylene glycol (PEG 1000, Merck, Carl Roth Ltd, Karlsruhe, Germany), and 100 μm sections were collected as previously described^5^. Neuropathological staging and disease classification were performed according to a previously published protocol^13^ by H.B. after AT8 immunostaining using a monoclonal antibody PHF-Tau (1:2000; Clone AT8; Pierce Biotechnology, Rockford, IL, USA (Thermo Scientific)) for recognition of phosphorylated tau protein in non-argyrophilic pretangle material and in argyrophilic NFTs of the Alzheimer type. Aβ deposition was staged using the monoclonal anti-Aβ antibody 4G8 (1:5000; Covance, Dedham, MA, USA) as described previously^37^. PART classification included cases with tau stages 1b-IV, Aβ phase 0. AD classification included cases with tau stages 1b-VI, Aβ phase ≥ 1. Subjects that met the criteria for “possible PART” (Aβ phases 1-2) were included with the remainder of AD subjects, given the presence of concomitant tau and Aβ pathology in these individuals^6^. In the present study, 20 cases displayed coincident argyrophilic grain disease (AGD), but care was taken to exclude other non-AD tauopathies, including progressive supranuclear palsy, Pick’s disease, and corticobasal degeneration. In addition, all cases were also immunostained and ^38^ staged for sporadic Parkinson’s disease (PD), as described elsewhere^38^. A total of 20 cases showed coincident α-synuclein-positive Lewy pathology.

**Table 1.**
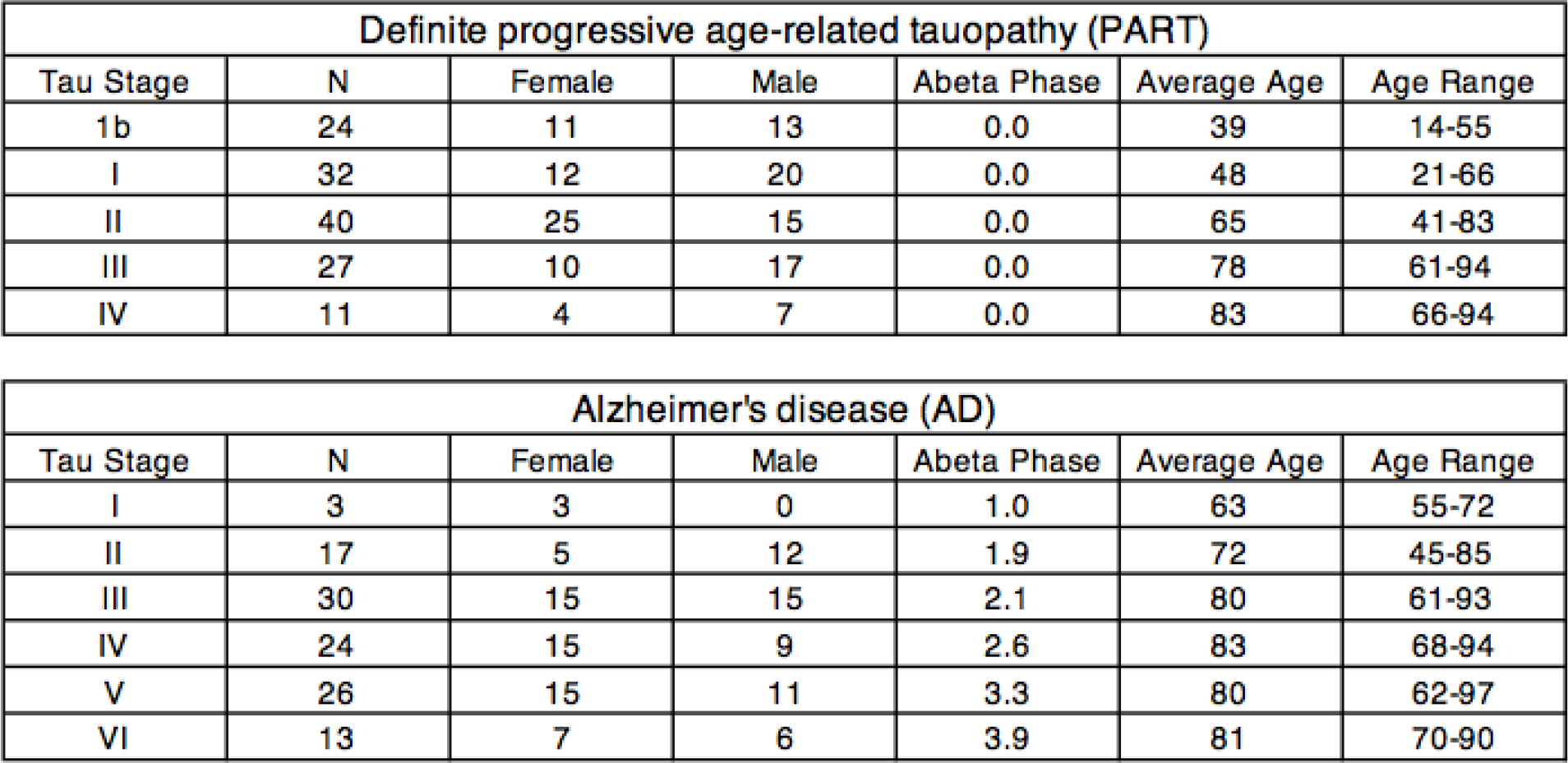
Summary of case samples.

From each case, including negative controls, 4 mm punch biopsies were collected from unstained sections of the locus coeruleus (LC); the transentorhinal cortex (TRE) and entorhinal cortex (EC) (two separate adjacent punches were taken from this combined region, termed TRE/EC, for seeding analyses); the first (superior) temporal gyrus (FTG); and the primary visual field (striate area, VC) with a punch biopsy tool (Kai Industries Co, Ltd. Japan) by K.D.T. To avoid cross contamination of seeding activity between individuals and regions, punch biopsy tools were used only once for each sample.

Samples were encoded and all subsequent preparation and seeding assays were performed in a blinded fashion. Tissue punches were stored in 1× TBS at 4°C until use. Samples were transferred to 100 μL of 1× TBS with protease inhibitors (Sigma Aldrich complete protease inhibitor, EDTA free), and water-bath sonicated in PCR tubes for 120 minutes under 50% power at 4°C (Qsonica Q700 power supply, 431MPX microplate horn, with chiller).

### Transduction of biosensor cell lines

Biosensor cells were plated at 25,000 cells per well in 96-well plates. After 18 hours, cells were transduced with human tissue homogenates as previously described^32–35^. Samples were added to Opti-MEM (Thermo Fisher Scientific) and incubated for 5 minutes (3.3 μL lysate with 6.7 μL of Opti-MEM per well). Lipofectamine was incubated with Opti-MEM (1.25 μL Lipofectamine with 8.75 μL Opti-MEM per well) for five minutes. Lipofectamine complexes were then mixed with samples and incubated for 20 minutes prior to addition to biosensor cells. Samples were assessed in triplicate. Cells were kept at 37°C in a humidified incubator for 48 hours, and subsequently dissociated with trypsin and prepared for analysis by flow cytometry.

### Flow cytometry and analysis of seeding activity

Biosensor cell lines were harvested with 0.05% trypsin, and quenched with media (DMEM + 50% FBS, 1% Pen/Strep, 1% Glutamax). Cells were spun at 500 × g and resuspended in 2% PFA in 1× PBS. Cells were subsequently spun and resuspended in flow buffer (HBSS + 1% FBS + 1mM EDTA) and stored for less than 24 hours prior to performing flow cytometry. All flow cytometry for biosensor cells treated with mouse-derived tissue was performed using a Miltenyi VYB flow cytometer. Flow cytometry for all human-derived samples was performed using a BD Biosciences LSR Fortessa. Flow cytometry data was analyzed as previously described^33^. Seeding activity was calculated as (percentage of FRET-positive cells)*(median fluorescence intensity), which was normalized to negative control samples (tau knockout mouse brain).

### Semiquantitative tau histopathology analysis

Individual microscopic slides from each case were staged for AD-associated lesions by
H. B. prior to decoding and analysis of the corresponding punch biopsies made from adjacent unstained tissue sections (S.K., T.T.). The LC, TRE/EC, FTG, and primary VC were assessed as follows: 0 = no detectable AT8-immunoreactivity, (+) = at least one AT8-immunopositive axon and/or cell soma, + = mild AT8-immunopositive pathology,
++ = moderate AT8-immunoreactive pathology, +++ = severe AT8-immunoreactive pathology.

### Statistical analyses

All samples collected by punch biopsy in Ulm were blinded to neuropathological stage prior to performing seeding assay analysis in the Diamond laboratory. All samples from an individual brain region were assessed in parallel with tau KO mouse brain samples. A stringent seeding threshold was set at 4 standard deviations (SD) above the average signal obtained from negative control tau KO mouse brain samples. Flow cytometry gating and analysis of seeding activity were completed prior to the decoding and interpretation of seeding results. All statistical analysis was performed using GraphPad Prism. Kruskal-Wallis one-way analysis of variance (ANOVA) with Dunn’s multiple comparisons test was performed to compare seeding between AD and PART subjects at NFT stages I-IV. Spearman r correlation was calculated for correlation of seeding activity between each brain region.

## Results

### Reproducible seeding activity in adjacent sections

We previously developed a protocol to compare seeding activity from fixed brain section punch biopsies in mice with AT8 staining in adjacent tissue sections^35^. To verify the reliability of this method in the human brain, we compared seeding activity in two adjacent 4 mm punch biopsies taken from the combined TRE/EC region in individual AD and putative PART patient fixed brain samples. We homogenized samples using sonication, and transduced lysate into previously described biosensor cells^32^. We then quantified tau seeding based on the degree of intracellular aggregation measured by FRET flow cytometry, relative to brain samples from tau knockout mice^32,35^. In these studies, we set a highly stringent threshold of 4 SD over background as a “positive” signal. In the present study, we observed good correlation between adjacent punch biopsy samples (n=247 cases, r = 0.9) (Fig. 1).

**Figure 1.**
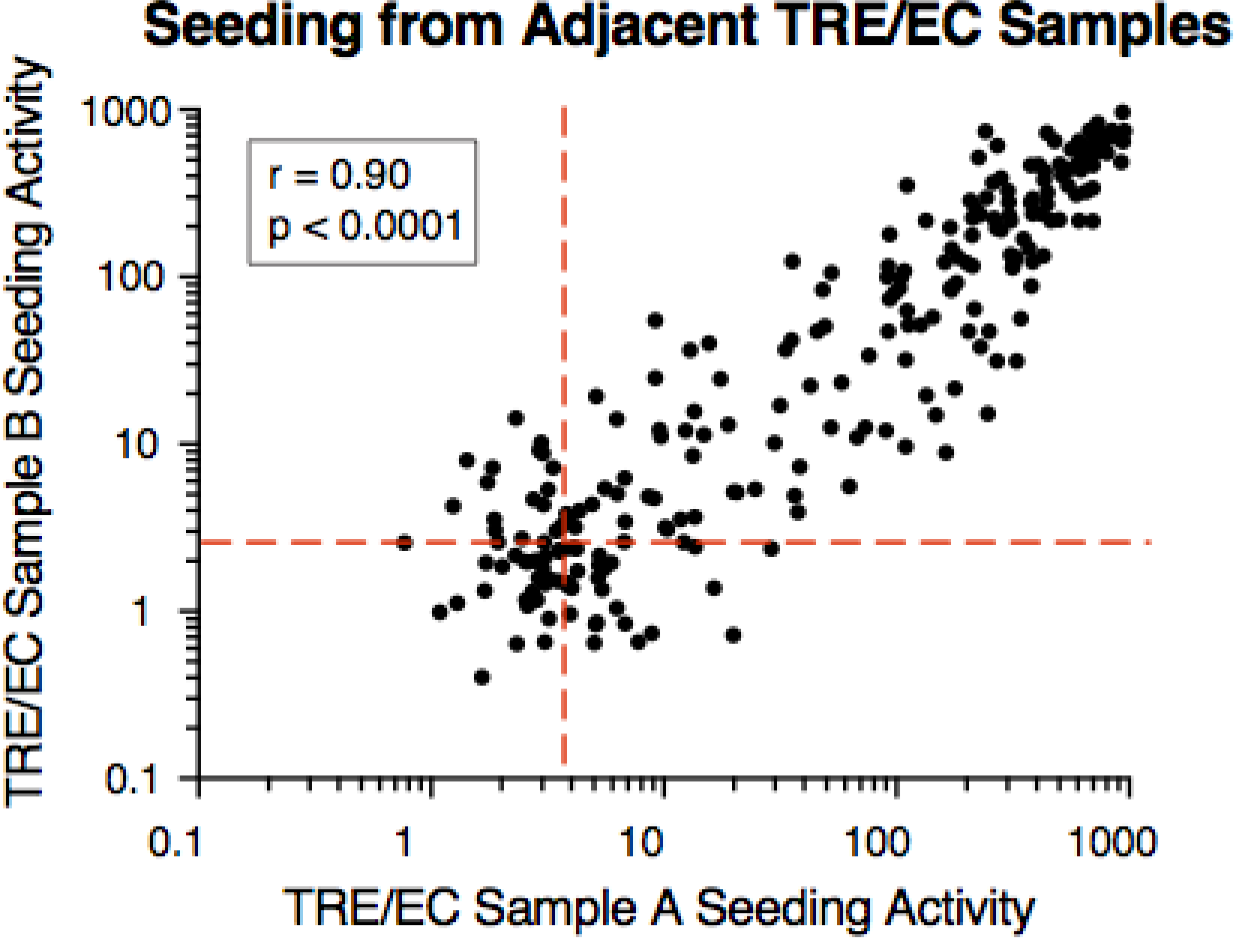
Tau seeding assay reliably detects tau aggregate pathology in formaldehyde-fixed tissue from NFT-staged cases. Seeding activity from adjacent punch biopsies correlated significantly with one another (n=247, P<0.0001). Two immediately adjacent punch biopsies were taken from the TRE/EC region and tested for phospho-tau seeding activity. Seeding activity correlated well between punches. Spearman r and p values are displayed on the graph.

### Seeding increases with higher NFT stages in AD and PART

Next we assessed seeding activity in a blinded fashion at progressive NFT AT8 pathology stages. We compared cases classified as the recently defined “definite progressive age-related tauopathy” (PART, NFT stages 1b-IV, Aβ phase 0) with AD cases (AD, NFT stages I-VI, Aβ phase ≥ 1). At NFT stage 1b, mild AT8 signal is present in the LC and in single or a few pyramidal cells in the TRE. In the LC biopsy punches, 8% of subjects displayed a small degree of seeding activity (Fig. 2A). In contrast, 29% of stage 1b subjects and over 50% of NFT stage I subjects displayed seeding activity in the TRE/EC punches (Fig. 2B). We detected robust seeding activity in the TRE/EC in over 90% of subjects at NFT stage II or higher (Fig. 2B). Seeding in the TRE/EC peaked by NFT stage IV and remained high in later disease stages (Fig. 2B). In the first temporal gyrus (FTG), where AT8 pathology in cortical projection neurons does not develop until NFT stage V, we detected seeding activity at NFT stage III in over 50% of individuals (Fig. 2C). Similarly, 33% of subjects exhibited seeding activity in the primary visual cortex (VC) as early as NFT stage III, although AT8 pathology in cortical nerve cells typically develops in this brain region only during the latest stages of AD (Fig. 2D). The seeding assay thus detects tau pathology prior to that which can be visualized by AT8 IHC in brain regions, such as the FTG and primary VC. Furthermore, our data are inconsistent with the LC as the “origin” of seeding in AD, as this region does not exhibit seeding activity consistently until NFT stages III-VI.

We detected no difference in seeding activity in the TRE/EC between AD and putative PART subjects at NFT stages I-IV (p>0.05, one-way ANOVA). As for AD, PART subjects also displayed positive seeding activity in brain regions, such as the first temporal gyrus and visual cortex in NFT stages II, III, and IV, despite the absence of AT8-positive NFT pathology. PART and AD exhibited similar overall patterns of progression and levels of tau seeding activity despite the differences in Aβ pathology.

**Figure 2.**
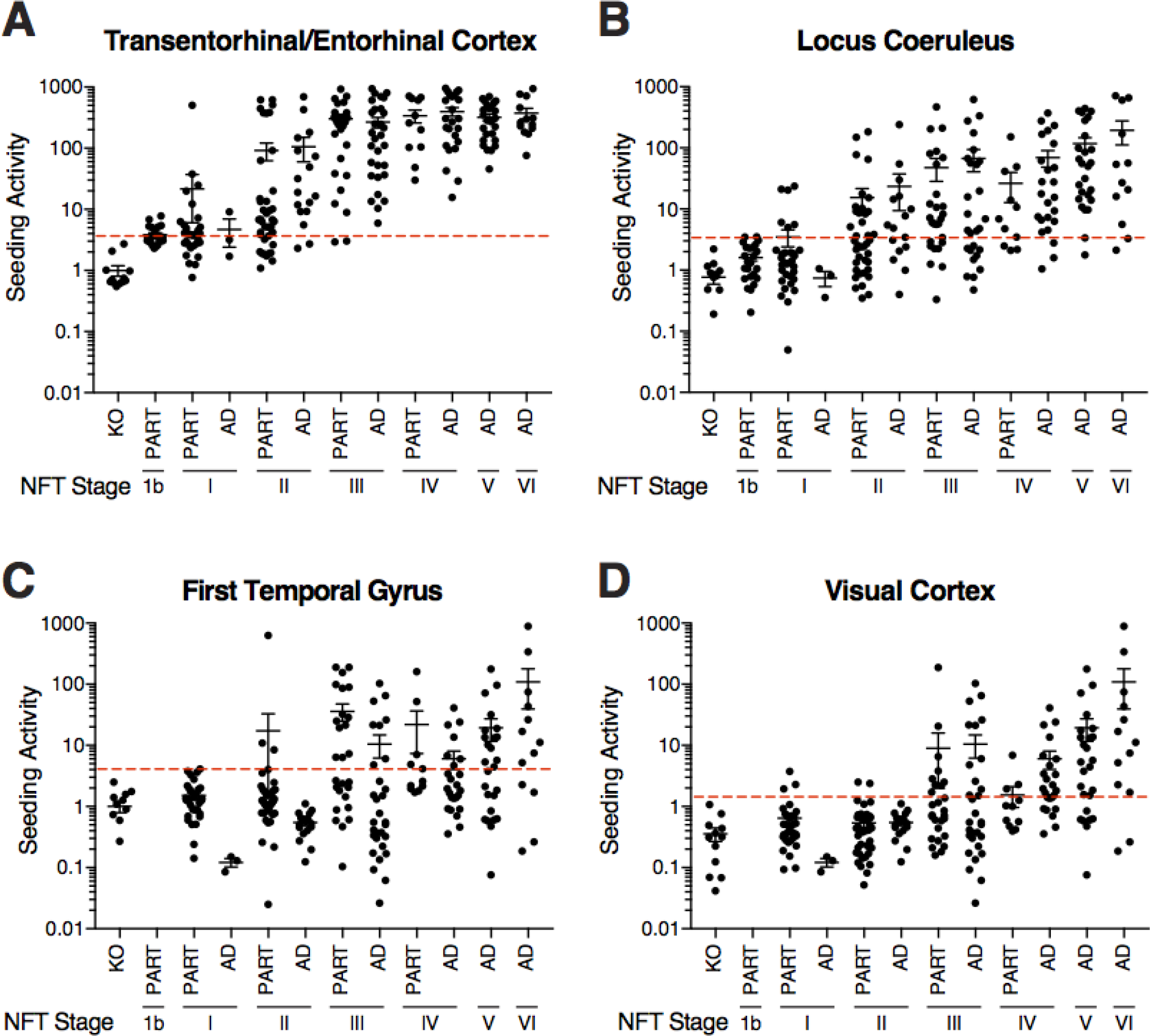
Tau seeding activity across brain regions. Tau seeding activity and NFT staging was performed blinded for each of four brain regions in n=247 subjects: TRE/EC, LC, FTG, and VC (striate area). For NFT stage 1b, samples were taken only from the LC and TRE/EC. (**A**) Seeding activity was first observed in the transentorhinal and entorhinal cortex (TRE/EC) at stage 1b, and increased several-fold at later NFT stages. Every individual examined showed positivity in this region by NFT stage IV. (**B**) Seeding in the LC was first detectable at NFT stage I in a small number of cases. Most samples exhibited tau seeding by NFT stage III. (**C**) Seeding activity in the FTG was observed in a limited number of cases by NFT stage II and increased at later stages. (**D**) The VC displayed positive seeding activity as early as NFT stage III, but approximately 15% of the individuals sampled did not show positivity even at NFT stage VI. KO=tau knockout mouse brain.

### Tau seeding vs. AT8 histopathology

NFT staging is performed by determining AT8 signal across multiple brain regions^5^, but direct comparison between IHC and tau seeding in AD required blinded analysis of AT8 signal in individual brain regions. Thus, we used AT8 to stain 100 μm brain sections immediately adjacent to those used for the seeding assay. We scored AT8-positive phospho-tau pathology on a semiquantitative scale (see methods section). We then plotted seeding activity against the assessment of AT8-positive staining in the LC, TRE/EC, FTG, and primary VC (Fig. 3A-D). We observed AT8-positive pathology in the absence of detectable seeding, particularly in the LC (Fig. 3A). We also observed tau seeding in the absence of tau AT8 pathology, most notably in the FTG and primary VC (Fig. 3C,D). However, the vast majority of samples with strong AT8-positive pathology also displayed robust seeding activity. These data were consistent with our prior observation that tau seeding anticipates AT8 immunostaining in cortical regions that typically score positive at late NFT stages^34^.

**Figure 3.**
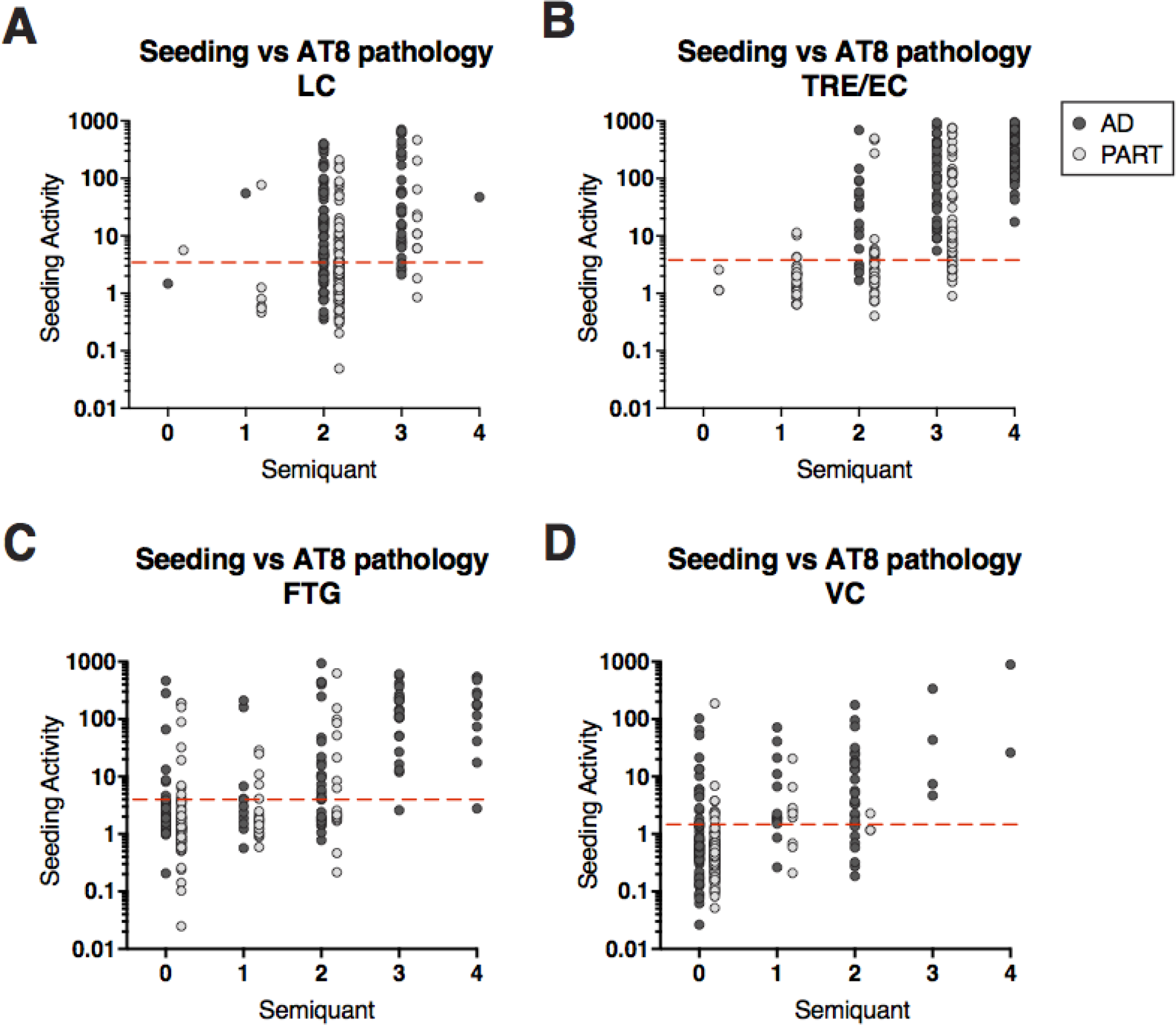
Tau seeding activity versus semiquantitative AT8 pathology. Seeding activity and AT8 histopathology were each performed blinded, and the results compared. (**A**) Subjects with a range of AT8 tau pathology (0=none,1=(+),2=+,3=++,4=+++) displayed robust seeding activity in the TRE/EC. Subjects with a higher degree of AT8 signal displayed higher levels of seeding activity. (**B**) Tau seeding activity in the LC was compared to AT8 signal. Subjects with mild to moderate tau AT8 pathology (levels 1-3) had a range of seeding activities, and a substantive number exhibited no seeding activity despite AT8 signal. (**C**) Seeding in the FTG was detectable prior to AT8 pathology in several AD and PART brain samples. (**D**) Seeding in the primary VC could be detected prior to AT8 signal in multiple AD and PART brain samples. Note PART subjects only spanned NFT stages 1b-IV.

### TRE/EC seeding precedes tau pathology in other brain regions

To further evaluate the pattern of progression of aggregated tau in different brain regions, we correlated tau seeding between the TRE/EC and other brain regions for individual subjects (Fig. 4). The TRE/EC exhibited seeding activity when other regions did not, consistent with the idea that the TRE/EC rather than the LC is the first region to develop pathogenic forms of tau. When seeding activity was compared between the LC, FTG, and VC, we observed a hierarchical pattern, with seeding developing in the LC and FTG without strong signal in the VC (Suppl. Fig. 2A-C).

**Figure 4.**
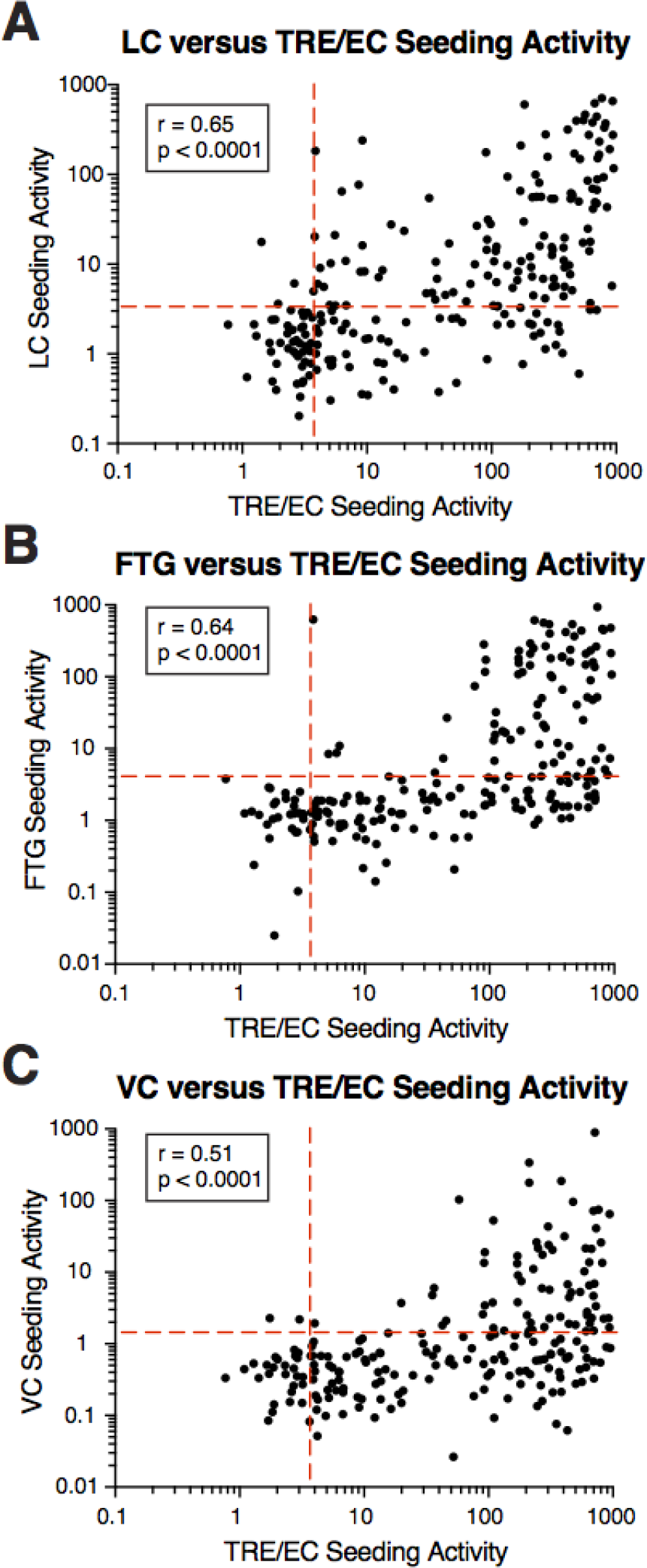
Correlations of tau seeding activity across brain regions. (**A**) Tau seeding activity was typically observed in the TRE/EC before seeding in the LC, and was higher in this brain region for the majority of subjects. Spearman r and p values are displayed on the graph. Seeding typically appeared first in the TRE/EC and at higher levels than in the FTG (**B**) or the VC (**C**). Spearman r and p values are displayed on the graph.

### Accumulation of tau seeding and AT8 pathology within subjects

To examine the progression of pathology across all brain regions, we created a heat map of tau seeding activity for each subject studied (Fig. 5). The TRE/EC reliably developed seeding first in AD and PART cases, and signal increased in all subjects at later NFT stages. Moreover, we observed a clear hierarchy within individual subjects, with the highest seeding typically appearing in the TRE/EC. We saw no consistent increase in seeding within the LC until NFT stage III. In contrast, we consistently observed early stage seeding and AT8 signal in the TRE/EC, and late stage increases in seeding and AT8 signal in the FTG. Several PART subjects also displayed seeding activity in the VC as early as NFT stage III, and we observed a clear gradient of seeding activity across individual subjects at increasing stages. Despite a similar pattern and degree of seeding in AD and PART subjects, a larger number of AD subjects had robust seeding in the VC at NFT stages III and IV (Fig. 5). However, this difference was not statistically significant at this number of cases (p>0.05, one-way ANOVA).

**Figure 5.**
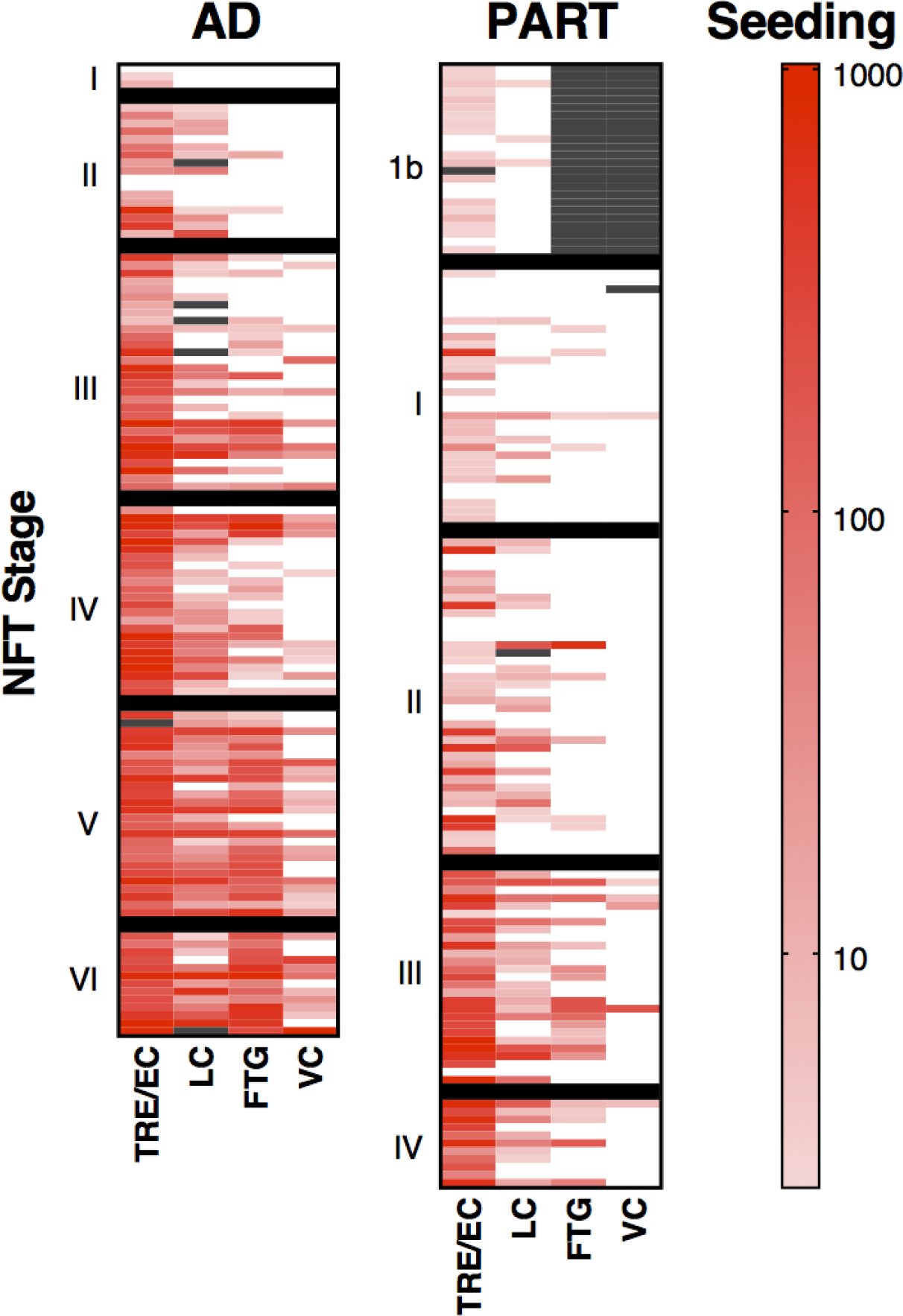
Seeding activity across multiple brain regions for individual AD and PART cases. Cases were categorized as probable AD vs. PART based on neuropathological criteria. Samples from each individual were directly compared across multiple brain regions. A continuous heat map of tau seeding activity (logarithmic scale) was plotted for each case and organized by staging and disease entity (AD, PART). AD subjects were arranged within each stage from low to high Aβ. Gray boxes represent unavailable samples. Seeding in the TRE/EC increased first and remained high for each disease stage. Subjects typically displayed less seeding in the LC, FTG and VC vs. the TRE/EC. The level of tau seeding in these secondary brain regions was higher at later NFT stages. Cases categorized as definite PART displayed a similar trend for the spatiotemporal progression of seeding activity when compared to AD. Grey boxes indicate absent samples.

## Discussion

To test fundamental ideas about AD and PART, we have used a highly sensitive and specific tau biosensor assay to measure seeding activity quantitatively in formalin-fixed brain tissues ~100 μm from adjacent sections staged by classical IHC. There has previously been uncertainty about the origin of AD pathology and whether it arises in the LC or the TRE/EC. Similarly, it remains unclear whether AD and PART constitute distinct neuropathological processes or are variants of the same disorder. Finally, it has not been definitively tested whether tau seeding activity in human brain anticipates subsequent NFT pathology, as would be predicted by the prion model.

Recent work proposes that AD and PART may be different diseases^6^. PART cases are defined to have minimal Aβ pathology or lack it entirely, and generally feature a relatively limited spread of tau pathology into cortical regions beyond the TRE/EC and hippocampus^6^. In this study we did not observe a pattern of tau histopathology in AD (i.e., with coincident Aβ pathology, n=113) that was clearly distinct from cases considered to represent definite PART (n=134), and seeding activity was similar in both conditions across the TRE/EC, LC, FTG, and VC. We observed a similar pattern of progression and similar seeding activity for both groups, across all neuropathological stages, despite different levels of Aβ deposition. This contrasts with a recent report of higher seeding activity in the presence of plaque pathology^39^. This may reflect that we sampled identical regions from the same fixed tissue block (separated by ~100 μm) instead of separate fresh and formalin-fixed tissues and evaluated a larger number of cases (n=247 vs. n=11). It remains unknown whether PART and AD might arise from distinct tau prion strains. Future work that examines the tau seed conformations (i.e., strains) present in these cases will help elucidate whether PART constitutes a separate disease entity^6^ or represents a type of prodromal AD^40^.

Despite early AT8 positive signal, we typically observed tau seeding activity in the LC only after it was already prominent in the TRE/EC, i.e., at later NFT stages (IV-VI). This is not consistent with the LC as the origin of tau seeding pathology. Instead, our data are consistent with the idea that tau seeds spread from the TRE/EC to the LC and then to more distant cortical regions, such as the FTG and subsequently the primary VC.

We have attempted to combine two orthogonal measures of pathology: classical IHC and a cell-based assay that depends on detection of bioactive tau seeding activity. Seeding and phospho-tau pathology did not uniformly correlate. For example, we observed AT8-positivity in the LC in the absence of seeding activity and seeding activity in the FTG and VC in the absence of clear NFT pathology. In this study we only examined seeding activity in brain regions that had been previously described to accumulate phospho-tau pathology by AT8 IHC, and thus were biased towards brain regions “classically” affected with NFTS. Indeed, other brain regions may also show seeding activity in the absence of AT8-positivity. To fully understand the relationship of seeding to pathology in AD, testing of multiple brain regions across NFT stages will be required. Interestingly, we note that in our prior study of seeding activity in fresh frozen tissue of patients with AD, we observed seeding activity in the cerebellum of 3/6 patients with late stage AD (a region that virtually never shows overt NFT pathology).

In testing various ideas about the origins and progression of AD and PART, this work is the first to combine a bioassay of tau seeding activity directly with classical histopathology on adjacent, formalin-fixed tissue sections. We observed no discernible differences between AD and PART with regard to AT8 staining at NFT stages I-IV. We also found no evidence to support the idea that early AT8 signal in the LC indicates that this region is the initial source of pathogenic seeding in AD. Instead, our data are consistent with the TRE/EC as the first site that develops tau seeding activity. Finally, we clearly observed that tau seeding activity anticipates detectable NFT pathology in the FTG and VC, consistent with the prion model of transcellular propagation of tau seeds as a driver of disease progression.

## Acknowledgements

S.K.K. and M.I.D. thank Nicolas Loof for providing guidance on flow cytometry techniques, and Matthew Brier for insightful discussion and statistical guidance. M.I.D., S.K.K., and T.L.T. were generously supported by the Rainwater Charitable Foundation and the NIH/NINDS. H.B. and K.D.T. thank the Goethe University Frankfurt (Braak Collection) and the Hans and Ilse Breuer Foundation (Frankfurt am Main, Germany) for generously supporting their research. This work was supported by Moody Foundation Flow Cytometry at the University of Texas Southwestern Medical Center.

---

**Supplemental Figure 1.**
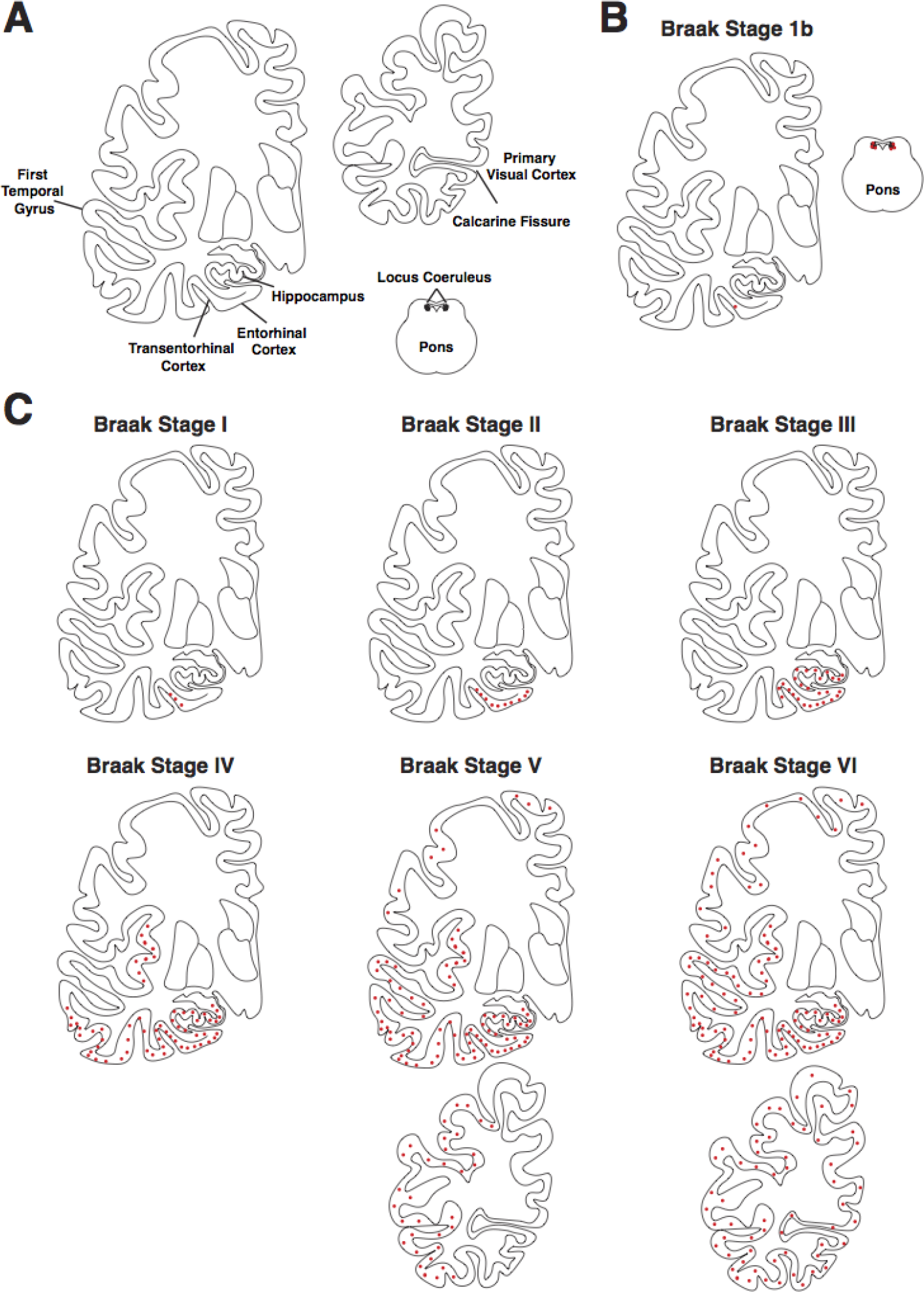
Summary of staging for AT8-positive tau pathology in AD. (**A**) Diagrams of brain regions of interest for AT8 phospho-tau pathology observed at different NFT stages. (**B**) Stage 1b is the earliest examined. Phospho-tau pathology is observed in the LC, and very limited pathology is present in the TRE. (**C**) NFT stages I-VI include increasing levels of phospho-tau pathology in specific brain regions. Stage I includes TRE pathology. This includes the EC by NFT stage II. Stage III includes pathology in the hippocampus. Stage IV includes pathology in the middle temporal gyrus and insula. Stage V includes pathology in additional cortical regions, including the first (superior) temporal gyrus. However, only NFT stage VI includes tau pathology in the primary visual neocortex (striate area). AT8 pathology is represented by red dots.

**Supplemental Figure 2.**
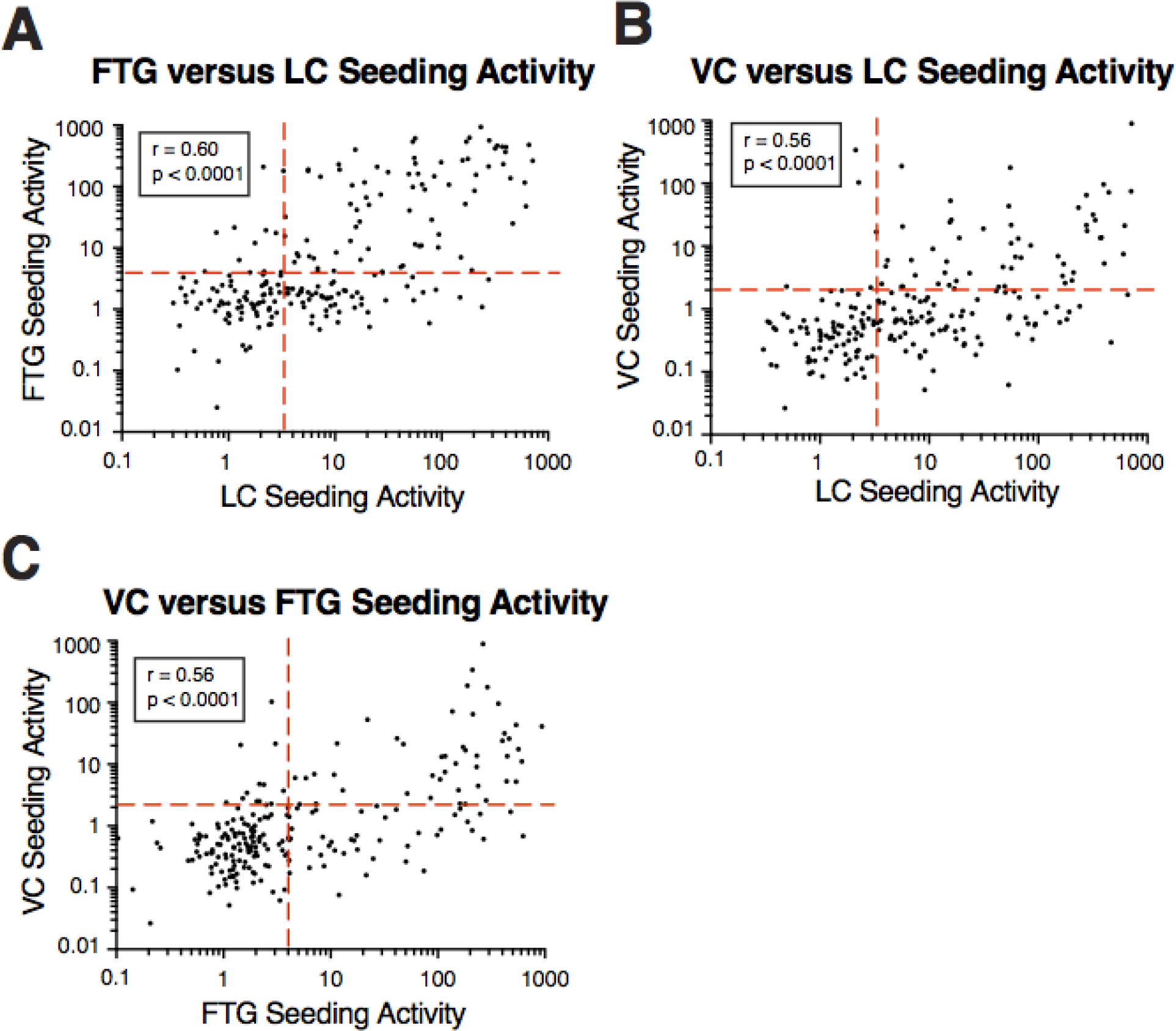
Correlation of tau seeding activity between the LC, FTG, and primary VC in cases with coincident AGD or Lewy body pathology. (**A**) Tau seeding in the LC was typically higher than in the FTG. However, several subjects displayed the opposite trend, with FTG showing seeding with no seeding in the LC. Spearman r and p values are displayed on the graph. (**B**) The LC typically displayed higher seeding than the VC. Spearman r and p values are displayed on the graph. (**C**) Seeding activity in the FTG was typically higher than in the primary VC. Spearman r and p values are displayed on the graph.

## Supplemental Table 1. Summary of AD-related neurofibrillary tangle (NFT) stages

**NFT stage I:** Gallyas silver staining reveals neurofibrillary lesions restricted to selected brainstem nuclei and the transentorhinal region (TRE).

**NFT stage II:** Tau pathology is present in the entorhinal cortex (EC) of the parahippocampal gyrus.

**NFT stage III:** Tau pathology in the CA1 sector of the hippocampal formation, and in neocortical regions of the temporal neocortex adjoining the TRE.

**NFT stages IV and V:** Increasingly prominent tau pathology in neocortical regions. The first temporal gyrus becomes involved at NFT stage V.

**NFT stage VI:** Tau pathology is present in neocortical areas, such as the primary visual cortex.

